# A naturally occurring Tat R52W variant in brain-derived HIV attenuates transcription and may contribute to reservoir persistence

**DOI:** 10.64898/2026.09.22.753475

**Authors:** Hongjie Chen, Nan Jiang, Yuyang Tang, Antoine Chaillon, Sara Gianella, David M. Margolis, Venkat R. Chirasani, Guochun Jiang

## Abstract

HIV-1 persistence under suppressive antiretroviral therapy (ART) remains a major barrier to eradication, yet the contribution of naturally occurring viral variation to transcriptional regulation is incompletely understood. Here, we identify an HIV-1 Tat variant (R52W) that attenuates viral transcription through the disruption of the Tat–TAR RNA interaction. Full-length HIV-1 genomes derived from the brain microglia of a “Last Gift” donor, together with peripheral viral sequences obtained during suppressive ART, revealed the presence of the R52W variant *in vivo*. The same R52W variant was found in another virally suppressed donor within the same cohort. Additional analysis of 3,440 clinical blood isolates from people with HIV identified R52W in 73 sequences (2.12%) of individuals, including three on ART and a therapy controller, and across diverse viral subtypes. Structural modeling demonstrated that R52W destabilizes Tat–TAR binding, while functional assays showed markedly reduced Tat-mediated transcriptional activity, which was restored upon tryptophan-to-arginine reversion (W52R) in Tat. These findings establish a direct molecular mechanism by which a naturally occurring HIV-1 variant modulates viral transcriptional output. Together, our results identify transcriptionally attenuated HIV-1 variants in treated individuals and suggest that modulation of Tat-dependent transcription may represent an underappreciated determinant of viral persistence under suppressive ART.

**Significance:** We identify a naturally occurring HIV-1 Tat variant (R52W) that attenuates viral transcription by disrupting Tat–TAR RNA interaction. This variant is detected in brain-derived and peripheral viral genomes and recurs across 3,440 clinical isolates from people with HIV, including multiple ART-suppressed individuals and a therapy controller. Functional and structural analyses demonstrate a direct mechanism for reduced Tat-mediated transcription. These findings reveal functional viral diversity under suppressive antiretroviral therapy and suggest that naturally occurring transcriptionally attenuated variants may contribute to HIV persistence in treated individuals.

## Introduction

Brain microglia represent a major latent reservoir of human immunodeficiency virus type 1 (HIV-1) in the central nervous system (CNS) of people with HIV (PWH) receiving suppressive antiretroviral therapy (ART)^1, 2^. As long-lived, self-renewing CNS-resident macrophages, microglia maintain stable integrated proviruses and may exhibit resistance to viral cytopathic effects and apoptosis^3^. Prior studies have identified adaptive changes of HIV within the CNS, including the emergence of compartmentalized, macrophage-tropic variants with distinct transcriptional properties^4, 5^. However, the availability of HIV-1 genomes derived specifically from brain microglia remains limited. A deeper understanding of viral genetic variation within these CNS-resident immune cells is therefore critical for elucidating the mechanisms of HIV persistence and informing strategies to address NeuroHIV^6^.

Efficient HIV-1 transcription requires interaction between the viral transactivator protein Tat and the transactivation response (TAR) RNA element, which together promote processive elongation by RNA polymerase II at the viral long terminal repeat (LTR) promoter ^7, 8, 9, 10^. TAR forms a conserved stem–loop structure containing a six-nucleotide apical loop and a three-nucleotide pyrimidine bulge that are essential for Tat binding and transcriptional activation^11^. The Tat protein comprises multiple functional domains, including a highly conserved arginine-rich motif (ARM; residues 49–59) within its basic domain, which mediates nuclear localization, high-affinity binding to TAR, and cellular uptake^12, 13, 14, 15, 16, 17^. Disruption of the Tat–TAR interaction impairs the recruitment of transcriptional elongation factors, leading to reduced viral transcription and contributing to the establishment and maintenance of HIV latency^18, 19^. Because this interaction is virus-specific, the Tat–TAR axis represents an attractive target for therapeutic modulation of HIV transcription without broadly affecting host transcriptional machinery^20, 21^.

Here, we analyzed two full-length HIV-1 genomes recovered from brain microglia isolated from a virally suppressed PWH enrolled in the “Last Gift” cohort, and identified a naturally occurring R52W substitution within the Tat ARM. This mutation markedly impaired Tat-mediated transcriptional activation, which was restored upon reversion to the wild-type residue. Our data suggest that the attenuation of Tat-mediated transcription is associated with viral persistence and identifies R52 in the Tat–TAR interface as a potential target for therapeutic strategy.

## Materials and Methods

### Brain microglia-derived HIV-1, viral sequence alignment, and global conservation and subtype distribution analysis of HIV Tat residue 52

Microglia-derived HIV-1 isolates were obtained from one PWH enrolled in the Last Gift Program (**Table 1**) ^22^. Replication-competent HIV-1 was recovered, and full-length viral genomes were previously sequenced^1^. Tat proteins were aligned using Clustal Omega and DNAMAN. The HIV-1 TAR secondary structure was predicted using mfold^23^, and compared with corresponding regions from the reference HIV-1 strain pNL4.3, HXB2, YU2, AD8, and JRCSF obtained from GenBank^24, 25, 26, 27, 28^.

**Table 1.** Participant information in this study.

| PID | Age | Sex | Plasma VL | Last VL days before death | Last CD4 count | Last ART | Stopped ART before death? |
| --- | --- | --- | --- | --- | --- | --- | --- |
| Pt1/Last Gift | 64 | Male | 68,007 | 1 | 100 | DRV/C/FTC/TAF | 111 days before death |
| Pt2/Last Gift | 75 | Male | <30 | 30 | 27 | N/A | No |
| Pt3 | N/A | Male | 199.53 | N/A | 757 | ZDV/3TC/ABC | N/A |
| Pt4 | 53 | Male | <40 | N/A | 440 | BAL | Therapy interruption |
| Pt5 | 29 | Male | <40 | N/A | 582 | N/A | N/A |

To evaluate the evolutionary conservation and prevalence of the Tat R52W substitution, HIV-1 Tat protein sequences were retrieved from the Los Alamos National Laboratory (LANL) HIV Sequence Database. Sequences representing multiple HIV-1 subtypes and circulating recombinant forms (CRFs) were downloaded using the “one sequence per patient” filter to minimize sampling bias and overrepresentation of closely related isolates. A total of 3,440 unique Tat sequences were included in the analysis. A total of 3,440 unique Tat sequences were included in the analysis.

Multiple sequence alignment was performed using Clustal Omega (v1.2.4). Residue numbering was assigned relative to the HIV-1 HXB2 reference sequence (accession K03455; sequence identifier B.FR.1983.HXB2LAIIIIBBRU.K03455). Amino acid conservation at Tat residue 52 was determined using a custom Python script that mapped HXB2 residue numbering onto the multiple sequence alignment and extracted the corresponding amino acid from each sequence. Amino acid frequencies were then calculated across the entire dataset and stratified by HIV-1 subtype and CRF where applicable.

To determine whether the R52W substitution was associated with specific viral lineages, metadata from the LANL HIV Sequence Database were used to classify isolates by HIV-1 subtype. The frequency of amino acid variants at Tat residue 52 was subsequently calculated for the major represented subtypes, including A, B, C, and D. To further assess evolutionary constraints within the Tat RNA-binding region, amino acid conservation across the arginine-rich motif (ARM) was visualized using sequence logo analysis^29^. This approach enabled qualitative assessment of residue conservation within the TAR RNA recognition domain and identification of positions exhibiting subtype-specific variability.

### Cell lines

TZM-bl cell line (a CXCR4-positive HeLa cell line engineered to express CD4 and CCR5 constitutively and firefly luciferase under the control of the HIV LTR was obtained from NIH AIDS Reagent Program and cultured in DMEM medium supplemented with 10% FBS. The Jurkat cell-derived 2D10 cell line of HIV latency was cultured in RPMI 1640 medium supplemented with 10% FBS. This cell line carries a single copy of the HIV-1 pNL4-3 provirus, which features full-length 5’ and 3’ LTRs but lacks most of the gag-pol region. The Nef gene is replaced with a green fluorescent protein (GFP) reporter gene, allowing for the monitoring of viral promoter activity.

### Full-length MG-OGV genome sequencing

Full-length HIV-1 genomes from outgrowth viruses (MG-OGVs) and Tat gene sequences were generated as previously described^1^. Briefly, cell-free viral RNA was extracted with Qiagen BioRobot EZ1 Workstation with EZ1 Virus Mini kit (Qiagen). Complementary DNA (cDNA) synthesis was performed in two separate reactions using primers B5R1 (for 5′ half-genome amplification) and R3B3R (for 3′ half-genome amplification). Near full-length HIV-1 genomes were then generated by limiting-dilution PCR followed by nested PCR amplification of overlapping 5′ and 3′ genome halves. Resulting amplicons were sequenced and assembled as previously described^1^. The full-length sequences of brain myeloid cell-derived HIV-1 strains, called BrMC2 and BrMC4, are provided in the Supplementary Materials.

### AlphaFold 3 and Tat-TAR interaction analyses

Variant-specific HIV-1 Tat–TAR complexes were generated using AlphaFold3^30^, as experimentally determined structures are unavailable for PWH-derived Tat and TAR sequence combinations. For each variant, the highest-ranking model was selected for subsequent molecular dynamics (MD) simulations.

Model confidence was evaluated using the AlphaFold3 confidence metrics, including predicted TM-score (pTM), interface predicted TM-score (iPTM), ranking score, predicted aligned error (PAE), and steric clash assessment. These confidence metrics are reported in **Supplementary Table 1**. All selected models were free of steric clashes and displayed consistent global folds and conserved Tat–TAR interaction topologies suitable for comparative MD simulations.

### Modeling of HIV Tat–TAR complexes

Seven HIV-1 Tat–TAR complexes corresponding to variants pNL4.3, AD8, BrMC2, BrMC4, YU-2, JRCSF, and HXB2 were obtained from the HIV Sequence Database^24, 25, 26, 27, 28^ (https://www.hiv.lanl.gov/). Sequences were aligned using Clustal Omega^31^ to identify conserved and variable residues at the Tat–TAR interface. The Tat-TAR complexes of eight variants were modeled using AlphaFold3^30^. Each AlphaFold3 model was assessed through predicted local distance difference test (pLDDT) scores and predicted aligned error (PAE) plots. The top-ranked models were selected for subsequent molecular dynamics (MD) simulations.

MD simulations were performed using GROMACS 2021.5^32^. The CHARMM36 all-atom force field^33^ was applied to both protein and RNA components. Each complex was placed in a triclinic box extending at least 1.0 nm from any atom to the box edge and solvated with TIP3P water molecules^34^. Sodium and chloride ions were added to neutralize the system and achieve an ionic strength of 0.15 M. Energy minimization was conducted using the steepest descent algorithm until the maximum force fell below 1000 kJ/mol/nm. The systems were equilibrated in two steps:

An NVT ensemble at 303.15 K for 100 ps using the V-rescale thermostat^35^ followed by an NPT ensemble at 1 bar for 100 ps using the Parrinello–Rahman barostat^36^. All covalent bonds involving hydrogen atoms were constrained using the LINCS algorithm^37^, allowing a 2-fs time step. Electrostatics were treated using the particle mesh Ewald (PME) method^38^ with a real-space cutoff of 12 Å, and van der Waals interactions were truncated at 12 Å. Production MD simulations were performed for 1 µs under constant temperature and pressure. Periodic boundary conditions were applied in all directions. The simulation trajectories were saved every 10 ps for structural and dynamic analyses.

### Trajectory and structural analyses

Trajectory analyses were carried out using built-in GROMACS utilities and custom Python scripts. Root mean square deviation (RMSD) analyses were performed to assess global conformational stability of the Tat–TAR complexes. Root mean square fluctuation (RMSF) profiles were calculated to identify flexible residues in Tat and regions of structural variability in TAR. Hydrogen bond analyses were performed using *gmx hbond* with donor–acceptor distance ≤ 3.5 Å and hydrogen–donor–acceptor angle ≥ 135°, yielding occupancy and time-dependent hydrogen-bond data between Tat and TAR. Secondary structure analyses were conducted using the DSSP algorithm ^39^ Implemented via *gmx do_dssp* to evaluate the stability of Tat helices during the simulation.

Visual inspection of representative structures and trajectories was performed using VMD 1.9.3 ^40^ and PyMOL 2.5 (Schrödinger, LLC). Comparative analyses among variants were used to correlate Tat secondary structure stability and H-bonding patterns with overall complex stability and conformational adaptability.

### HIV-1 Tat plasmids and plasmid transfection

HIV-1 Tat plasmids were prepared in VectorBuilder and Tat sequences were confirmed by sequencing. For the TZM-bl cell line, plasmid transfection was performed using the Lipofectamine 3000 Transfection Reagent (Invitrogen) following the manufacturer’s protocol. For the 2D10 cell line, transfection with Tat and TAR-2 plasmids was conducted using the SE Cell Line 4D-Nucleofector® X Kit (Lonza). Briefly, after washing with PBS, 1×10⁶ 2D10 cells were resuspended in 100 μL of a supplement/solution buffer mixture (mixed at a ratio of 1:4.5) containing the plasmids. The cell suspension was then transferred into a single cuvette and transfected using the 4D-Nucleofector® X Unit with program CK116.

### Luciferase reporter assay

TZM-bl cells (1×10⁴ cells per well) were seeded in a 96-well black plate one day before the experiment. Then, the cells were transfected with plasmids. Forty-eight hours after transfection, the cells were washed with PBS and incubated with 20 μL per well of Passive Reporter Lysis Buffer (Promega). Following overnight storage at –80 °C, 100 μL per well of Luciferase Assay Buffer (Promega) was added, and the luminescence signal was measured using a BioTek Synergy H1 Multimode Reader. Each experiment was performed with at least three independent replicates.

### Generation of HIV and HIV infection in human primary microglia

The pNL4-3-Δ6-GFP virus was obtained from Siliciano’s laboratory, which has a non-functional env gene that was replaced with a GFP reporter ^41^. The human 293T cell line was transiently transfected with the pNL4.3-Δ6-GFP plasmids along with PAX2 and VSVG with Lipofectamine 3000 (Invitrogen, L3000015). Viruses were harvested 48 hours post-transfection, filtered, concentrated, and titrated in 293T cells. HIV was used to infect microglial cells (Celprogen *via* shipment of frozen ampule, isolated from human brain cortex) via centrifugal inoculation at 1200 x g for 2 hours. The medium was changed 24 hours post-infection, and the cells were cultured for 48 hours. The GFP-positive cells were sorted for successfully infected cells, which were further cultured until undetectable for the latency study. Continuous culture after selection found that the basal HIV expression remained low (<2%), confirming that HIV was latently infected in microglial cells.

### Real-time polymerase chain reaction

Total RNA was extracted using the RNeasy Kit (Qiagen) and subsequently treated with DNase I (Invitrogen) to remove genomic DNA^1^. First-strand cDNA was synthesized with Superscript IV reverse transcriptase (Invitrogen) and random primers. Quantitative real-time PCR was performed using TaqMan chemistry on an ABI QuantStudio 5 system (Applied Biosystems) with gene-specific probe sets purchased from ThermoFisher Scientific. For TAR2, similar primers were used for SYBR PCR with a slight change: HIV-2 TAR-5 FCCCGCTTCTTGCATTGTATT and HIV-2 TAR-3 RACACTTAACTTGCTTCTAACTGG^42^. qPCR condition: 95°C 30 sec, 52°C 60 sec for 40 cycles. For HIV-1, the long LTR region was amplified for RT-qPCR, thereby representing HIV transcriptional elongation^19^. The expression levels were normalized to the host gene SDHA controls.

### Flow cytometry

HIV-1 expression and/or gene transcription were assessed by measuring GFP expression in 2D10 cell lines *via* flow cytometry, as we reported previously^43^. A viability dye (Invitrogen Aqua Live/Dead Fixable Dead Cell Stain) was used to distinguish live cells from dead cells. Data analysis was performed using FlowJo software.

### Whole cell, nuclear, and cytoplasmic protein Western blotting

Whole-cell lysates were prepared using RIPA lysis buffer supplemented with Protease/Phosphatase Inhibitor Cocktail (Cell Signaling Technology, Cat#: 5872). Nuclear and cytoplasmic proteins were extracted from TZM-bl cells 48 hours after transfection with Tat plasmids, using the NE-PER Nuclear and Cytoplasmic Extraction Reagents Kit (Thermo Scientific, Cat. # 78835). Protein levels were detected by anti-flag antibodies (F3165, ANTI-FLAG M2 antibody, Millipore-Sigma) and quantified by densitometric analysis using Image Lab Software (Bio-Rad).

### Quantification and statistical analysis

Specific statistical tests used are indicated in the figure legends, with significance levels denoted as follows: ∗*p* < 0.05, ∗∗*p* < 0.01, and ∗∗∗*p* < 0.001. All statistical analyses were performed using GraphPad Prism software, version 9.3 (GraphPad Software, USA). Data are presented as mean ± SEM where appropriate. Group comparisons were carried out using two-tailed Student’s *t*-tests for two-group comparisons, or one-way ANOVA for multiple-group comparisons. A *p*-value of less than 0.05 was considered statistically significant.

## Results

### Recovery and sequencing of outgrowth microglia-derived HIV-1 from brain tissue of PWH enrolled in the “Last Gift” Program

Brain tissue specimens were obtained via a rapid research autopsy from donors enrolled in the Last Gift cohort who underwent ART interruption and experienced viral rebound (**Table 1**). Following overnight shipment, tissues were immediately processed into single-cell suspensions using mechanical dissociation and enzymatic digestion, as previously described^1^. Microglia were isolated by positive selection for CD11b following depletion of CD3⁺ T cells, yielding a highly enriched microglial population (**Fig. 1A**). More than 95% of CD3⁻/CD11b⁺ cells expressed the microglia-specific marker TMEM119^1^, confirming their microglial identity. These highly purified microglia were used for subsequent virological and genomic analyses.

**Figure 1.**
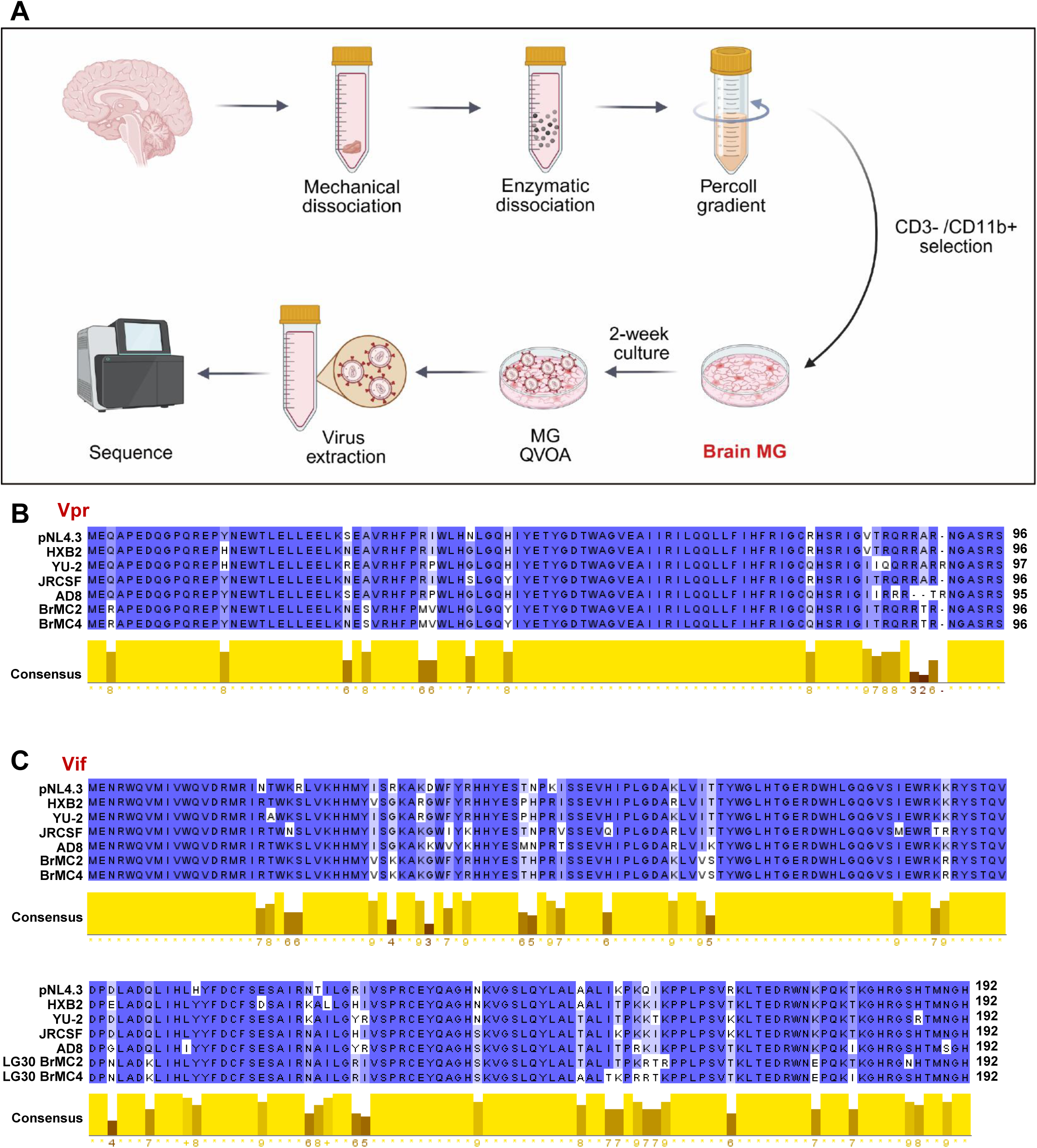
Genetic characterization of replication-competent HIV-1 isolated from brain microglia of PWH. **(A)** Schematic overview of the experimental workflow for the isolation of brain microglia (MG) from PWH and the subsequent HIV-1 sequencing. **(B and C)** Amino acid sequence alignments of HIV-1 Vpr (**B**) and Vif (**C**) comparing viral sequences isolated from brain microglia (BrMC2 and BrMC4) with commonly used reference HIV-1 strains (pNL4.3, HXB2, YU-2, JRCSF, and AD8).

Two weeks post-selection, the cells were amplified using the standard viral outgrowth assay (VOA) in co-culture with CD8-depleted PBMCs. Culture supernatants from purified microglia were harvested, and viral extracts were prepared for sequencing following limited dilutions of cDNA, as schematically summarized in **Fig. 1A**. The same approach was used to sequence microglia-derived HIV-1, as we previously reported^1^; in which only two clones from ∼10 VOA wells with relatively high viral load could be sequenced at the University of Pennsylvania. One isolate originated from the parietal cortex, and the other from the hippocampus of the PWH brain. These two isolates were designated BrMC2 (hippocampus) and BrMC4 (parietal cortex).

### Distinct Sequence Landscape of Tat and TAR of HIV-1 Derived from the Brain Microglia

Using these HIV genetic sequences, we performed multiple sequence alignments of key HIV-1 proteins—Vpr (**Fig. 1B**), Vif (**Fig. 1C**), Env, Vpu **(Supplementary Fig. 1)**, Gag, and Nef **(Supplementary Fig. 2)**. For comparison, the analysis included well-characterized T cell–derived strains (pNL-4.3 ^24^ and HXB2 ^25^) as well as brain-derived isolates (YU-2^26^, JRCSF ^27^, and AD8 ^28^). Sequence alignment showed that both Vpr and Vif were highly conserved across the isolates, with limited variation compared with pNL4.3, HXB2, or JR-CSF HIV-1. By contrast, Env, Gag, Nef, and Vpu were notably more divergent across these isolates, consistent with their known involvement in CNS infection and microglial tropism **(Supplementary Figs. 1-2)**.

Although overall conservation varied across proteins, the functional implications are most apparent when focusing on regulatory elements involved in viral transcription. Among these, the interaction between Tat and TAR has been proposed as a critical determinant of HIV-1 transcription efficiency^9, 10^. The Tat-TAR interaction recruits and stabilizes transcriptional machinery, thereby enhancing transcriptional initiation and elongation. In the presence of Tat, LTR-mediated transcriptional activity can be enhanced tens to hundreds of fold^8^, whereas viral replication drops to nearly undetectable levels in the absence of Tat. Given its pivotal role, we focused our analysis on sequence variations in Tat and their potential impact on this essential interaction.

Consistent with brain-derived HIV-1 strains YU-2, JRCSF, and AD8, which encode full-length 101-aa Tat proteins, but distinct from the truncated 86-aa Tat proteins encoded by pNL4.3 and HXB2, both microglia-associated HIV isolates encoded 101-aa Tat proteins with multiple amino acid substitutions across their functional domains^44^ (**Fig. 2A**). Among these, the Glutamine-rich domains (GRD) showed the highest degree of sequence variability, consistent with a previous report^8^. In contrast, the Arginine-rich region (ARM), encompassing the highly conserved motif 49RKKRRQRRR57 critical for TAR RNA binding as well as Tat secretion and uptake, was largely conserved across the analyzed strains.

**Figure 2.**
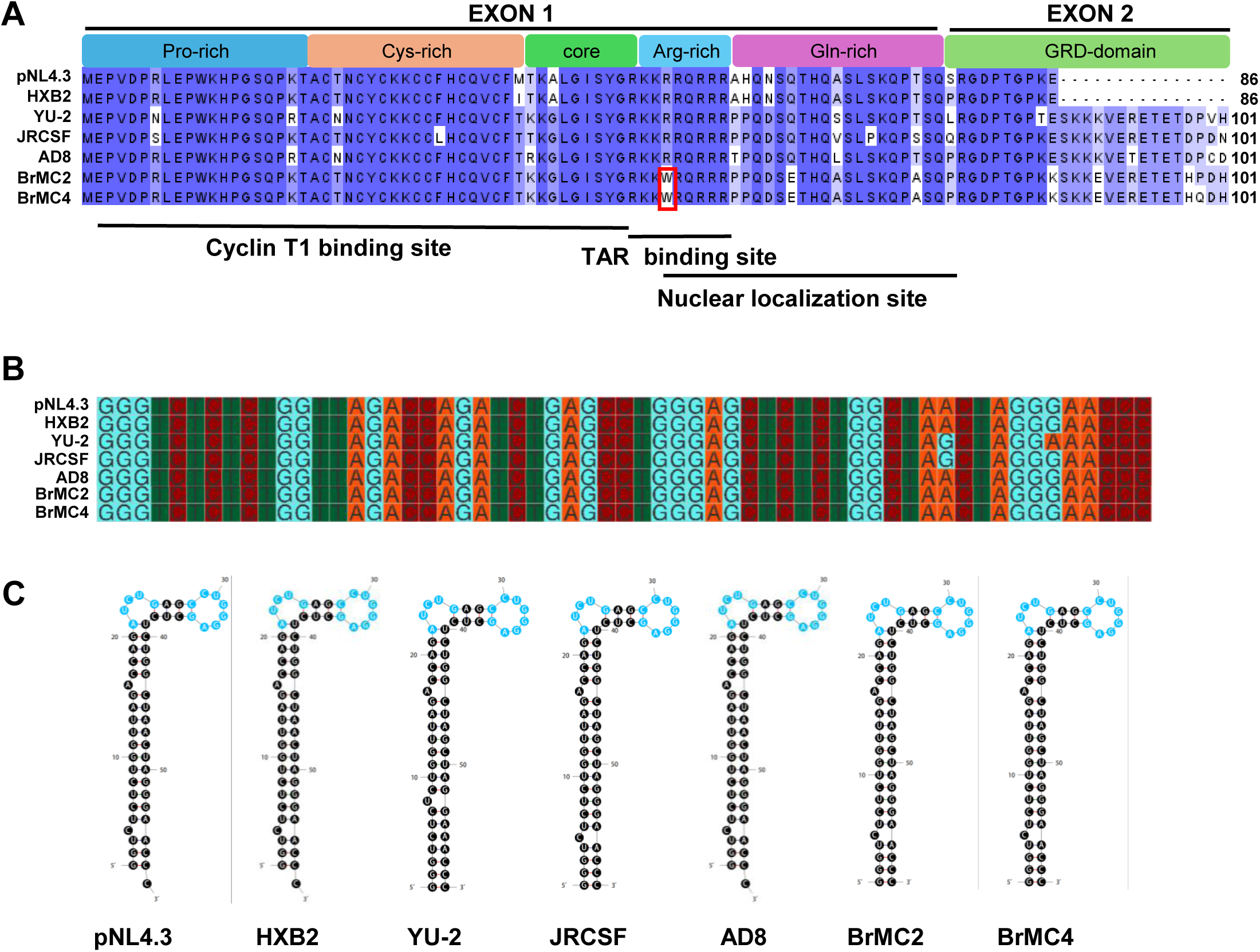
Sequence and structural analysis of HIV-1 Tat and TAR from brain microglia isolates. **(A)** Amino acid sequence alignment of the HIV-1 Tat proteins. The alignment compares Tat sequences from brain-derived isolates with reference HIV-1 strains pNL4.3, HXB2, YU-2, JRCSF, and AD8. **(B)** Nucleotide sequence alignment of the HIV-1 TAR RNA elements. **(C)** Predicted secondary structures of the TAR RNA from the corresponding viral isolates and reference strains.

Notably, the brain-derived isolates BrMC2 and BrMC4 harbored an R52W mutation within the ARM. Given the central role of this motif in TAR recognition, this substitution may alter Tat–RNA interactions and downstream transactivation efficiency. Additionally, amino acid changes were observed in the proline-rich, Core domain, and glutamine-rich domains relative to T-tropic HIV-1 and other CNS-derived HIV-1 strains. These regions have been implicated in T1 binding, TAR binding, nuclear localization, and transcriptional activation^8, 44, 45^.

In contrast to the observed variability in Tat, nucleotide alignment of the HIV-1 TAR RNA element demonstrated a high degree of sequence conservation across all isolates, including BrMC2 and BrMC4 (**Fig. 2B**). Secondary structure predictions showed that key structural features, particularly the bulge and apical loop regions essential for Tat binding and transactivation, were well preserved among all sequences (**Fig. 2C**). Together, these findings suggest conservation of TAR RNA structural integrity despite substantial sequence variation in microglia-derived Tat proteins.

### HIV-1 Tat-TAR interaction impairment in BrMC2 and BrMC4 HIV strains

To examine how Tat sequence variation influences the structural dynamics of microglia-derived HIV-1 Tat–TAR complexes, we modeled seven representative viral variants (pNL4.3, AD8, BrMC2, BrMC4, YU-2, JRCSF, and HXB2) using AlphaFold3^46^ (**Fig. 3A**). Across all variants, predicted global confidence ( pTM-score, i.e., pTM= ∼0.32–0.38) and interface confidence (ipTM= ∼0.15–0.23) were comparable, and no steric clashes were detected. Although minor variation in interface metrics and predicted aligned error (PAE) was observed, these differences did not alter the predicted Tat–TAR interaction topology (**Supplementary Table 1**). In all models, the Tat basic domain engaged the TAR RNA hairpin in a consistent orientation, supporting a conserved binding mode across viral variants. Collectively, the uniformity of model confidence metrics supports the robustness of comparative structural interpretations and indicates that observed differences are driven primarily by sequence-dependent effects rather than model uncertainty.

**Figure 3.**
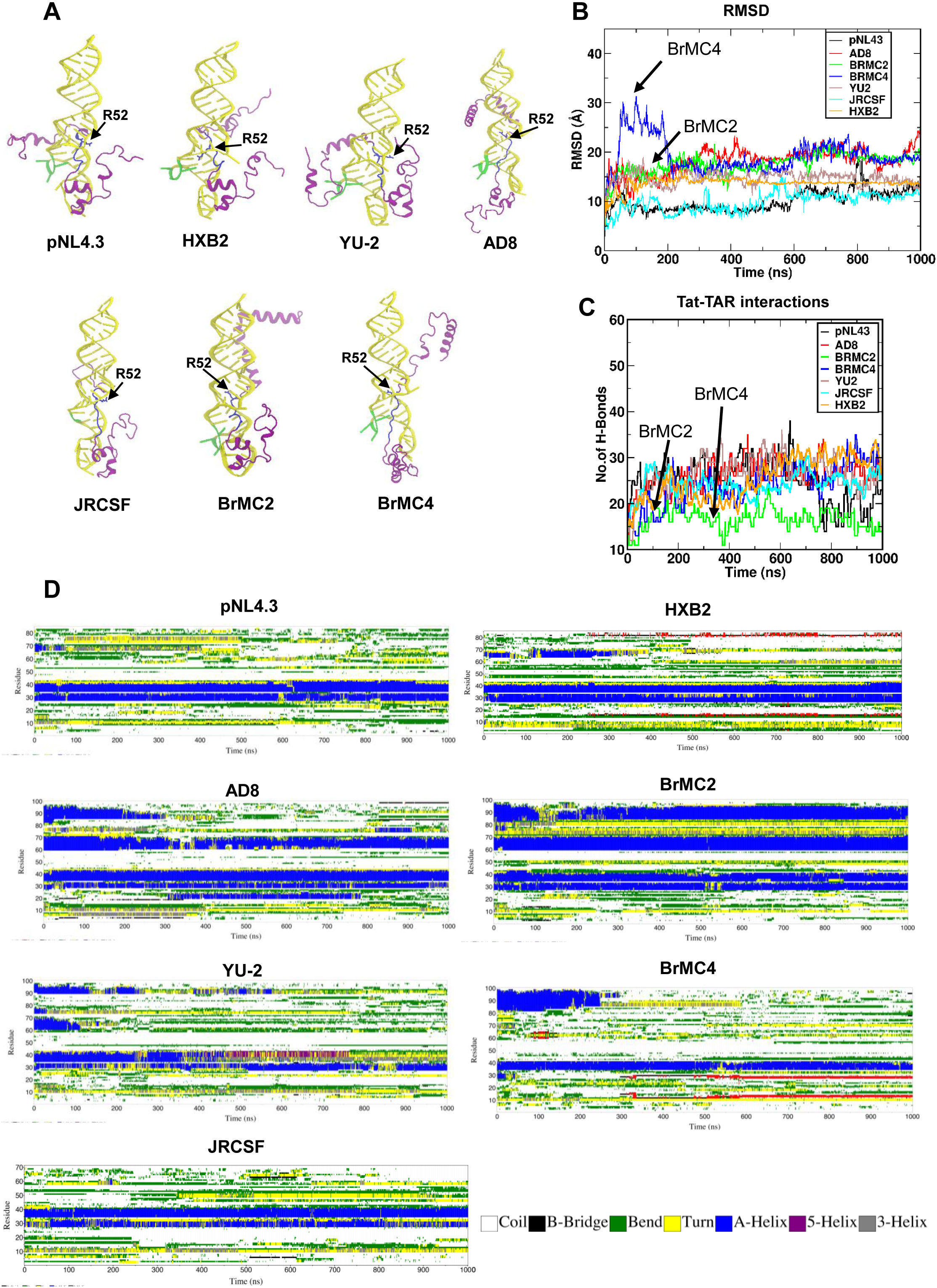
Structural modeling and molecular dynamics characterization of HIV Tat–TAR complexes across diverse viral variants. **(A)** AlphaFold3-predicted structures of HIV Tat–TAR complexes. Predicted Tat–TAR complex structures for seven HIV-1 variants (pNL4.3, AD8, BrMC2, BrMC4, YU-2, JR-CSF, and HXB2) generated using AlphaFold3. Tat proteins are shown in purple and TAR RNA hairpins in yellow. The arginine-rich motif (ARM) domain of Tat (residues ∼49–57) is highlighted in blue, with Tat residue R52 displayed in blue licorice representation. The TAR RNA bulge (nucleotides 23–25), critical for Tat binding, is highlighted in green. All models exhibit conserved Tat–TAR binding topologies, with variant-specific conformational differences in Tat influencing RNA recognition and complex stability. **(B and C)** Root mean square deviation (RMSD) profiles and intermolecular hydrogen bond counts were monitored over 1000 ns molecular dynamics simulations. The pNL4.3 Tat–TAR complex demonstrated the highest conformational stability, reflected by the lowest RMSD and persistent hydrogen bonding. In contrast, AD8, BrMC2, and BrMC4 exhibited higher RMSD values, indicating increased structural flexibility. BrMC2 formed the fewest stable hydrogen bonds, consistent with its reduced complex stability. **(D)** Time-resolved secondary structure analysis of Tat proteins in all Tat–TAR complexes over 1000 ns simulations. AD8 and BrMC2 retained prominent α-helical regions throughout the trajectories, indicating enhanced local stability. Secondary structure elements are color-coded as follows: α-helix (blue), β-bridge (black), bend (green), turn (yellow), π-helix (red), and 3₁₀-helix (gray).

To evaluate the reliability of the AlphaFold3-derived starting structures, we examined the prediction confidence metrics for all eight Tat–TAR complexes (**Supplementary Table 1**). The highest-ranked model was selected for each variant based on the AlphaFold3 ranking score. Across all systems, pTM values ranged from 0.31 to 0.41 (mean 0.35 ± 0.04), whereas iPTM values ranged from 0.15 to 0.25 (mean 0.20 ± 0.04). Importantly, all selected models were free of steric clashes and consistently reproduced the canonical Tat–TAR binding architecture, including positioning of the Tat ARM motif adjacent to the TAR bulge. Although the absolute interface confidence values were modest, likely reflecting the intrinsic flexibility of the Tat protein and current limitations in confidence estimation for protein–RNA complexes, the models provided a consistent structural framework for subsequent comparative molecular dynamics analyses.

To interrogate the dynamic stability of these complexes, we performed 1 µs MD simulations and analyzed root-mean-square deviation (RMSD) and intermolecular hydrogen bonding (**Fig. 3B**). RMSD profiles revealed distinct stability patterns across variants. pNL4.3 and JRCSF complexes exhibited the lowest RMSD values (∼10 Å), consistent with high structural stability. HXB2 and YU-2 complexes displayed intermediate RMSD stability (∼15 Å) (**Fig. 3B and Supplementary Fig. 3**). In contrast, BrMC2 and AD8 maintained persistently elevated RMSD values (∼20 Å), indicative of increased conformational plasticity. BrMC4 exhibited an early conformational transition within the first 200 ns followed by partial stabilization at elevated RMSD (∼20 Å), suggesting an unstable binding landscape.

Consistent with these dynamics, hydrogen bond analysis revealed marked variability in Tat–TAR interfacial stability (**Fig. 3C**). pNL4.3 complex initiated with∼30 Tat–TAR hydrogen bonds, which gradually decreased to ∼20, reflecting a dynamic yet stable binding interface. The JRCSF complex maintained ∼25 persistent hydrogen bonds, whereas HXB2, AD8, BrMC4, and YU-2 exhibited an average of ∼30 hydrogen bonds over the course of the simulation. By contrast, BrMC2 exhibited a reduced population of stable hydrogen bonds despite comparable RMSD values, indicating weaker, less persistent interfacial contacts.

Secondary structure analysis of Tat over the simulation trajectory further revealed variant-dependent conformational stability (**Fig. 3D**). Most variants (HXB2, AD8, BrMC2, JRCSF, and pNL4.3) maintained a stable α-helix–turn–helix motif between residues 30–40, indicative of a structurally constrained core. In contrast, BrMC4 and YU-2 Tat proteins exhibited loss of one helical element in this region, consistent with reduced structural rigidity. Additional α-helical elements formation in the C-terminal region was variably observed: BrMC2 displayed two persistent helices spanning residues 60–70 and 85–95, while AD8 maintained a stable helix at residues 60–70 and a transient C-terminal helix around residues 85–95. Similar transient helices were noted in BrMC4 and YU-2.

Notably, despite the preservation of local secondary-structure elements in BrMC2 and AD8, their high RMSD values indicate substantial global rearrangement and increased flexibility of the Tat–TAR interface. Together, these data indicate that sequence variation in CNS-derived HIV-1, particularly BrMC4 and BrMC2, destabilizes Tat–TAR complex formation through altered conformational dynamics and reduced or less persistent interfacial hydrogen bonding. These effects may contribute to variant-specific modulation of transcriptional efficiency and HIV gene regulation in brain-resident microglia.

### BrMC2 and BrMC4 Tat Variants have attenuated activities in the transactivation of HIV

To determine whether sequence variation in microglia-derived Tat proteins affects transcriptional activity, Tat genes from the indicated HIV-1 strains were cloned into pcDNA3.0 expression vectors and evaluated in two independent HIV reporter systems. We then transfected the TZM-bl reporter cell line with the corresponding Tat expression plasmids. A significant decrease in relative luminescence units (RLU) was observed in TZM-bl cells expressing the BrMC2 and BrMC4 Tat variants compared to T-tropic HIV-1 pNL4.3 or HXB2 (Fig. 4A).

**Figure 4.**
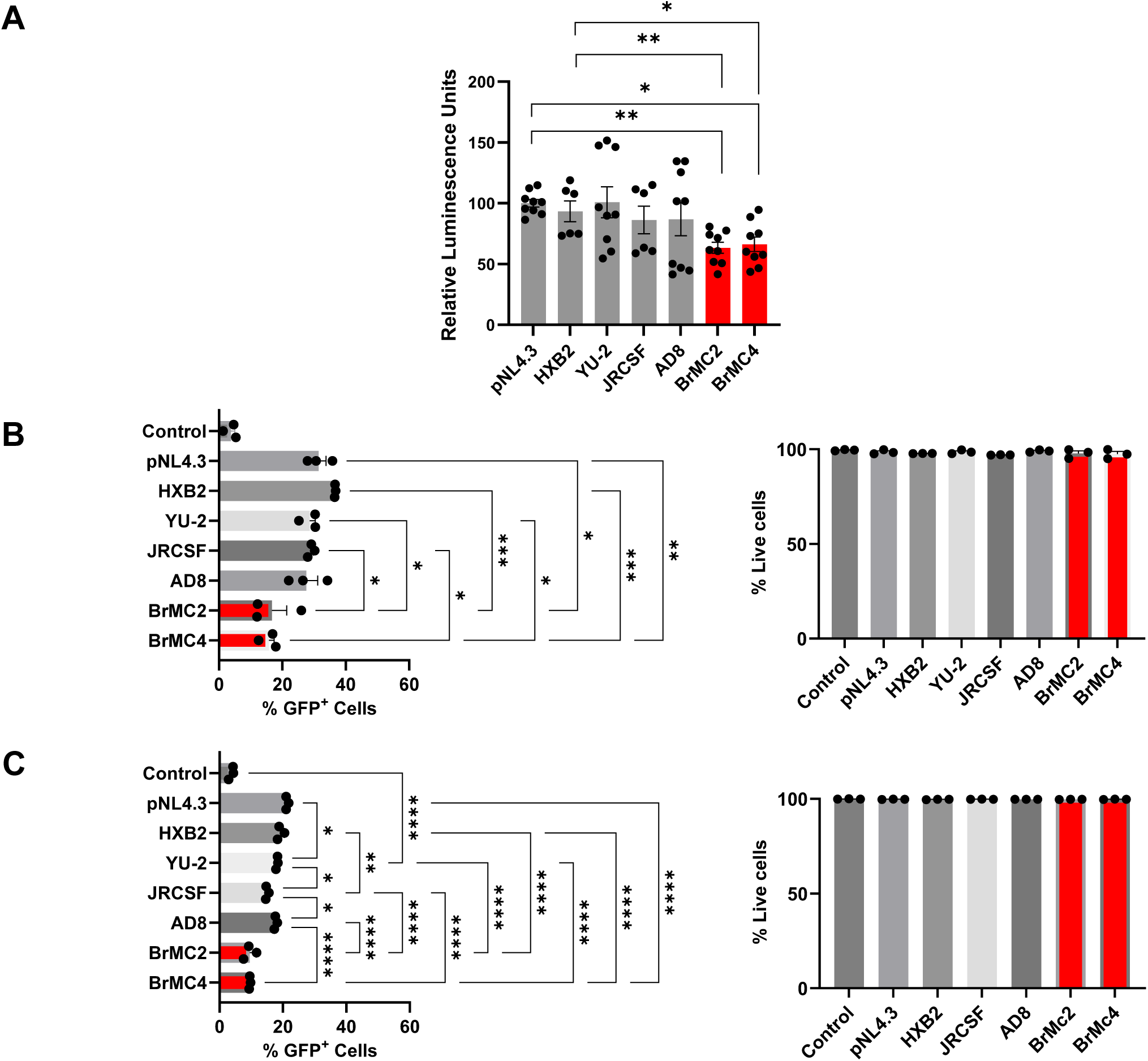
The R52W mutation impairs Tat-mediated transactivation in brain-derived HIV-1 isolates. **(A)** Luciferase assay of 2D10 cells 48 hours post-transfection with different HIV-1 Tat plasmids (1, 2.5, or 5 µg). *, *p*<0.5; **, *p*<0.01; and ***, *p*<0.001 with one-way ANOVA for multiple-group comparisons **(B-C)** GFP flow cytometry of 2D10 cells (B) and microglia (C) 7 days post-transfection with 2 µg HIV-1 Tat plasmids. The viability of the cells was also determined using flow cytometry.

This defect was independently validated in the Jurkat-derived latency model 2D10 after transfection with BrMC2 and BrMC4 plasmids, indicating impaired Tat-mediated transactivation (∼55% reduction for BrMC2 and ∼57% reduction for BrMC4) (**Fig. 4B**). Similar defects were observed in latently infected microglial cells, where BrMC2 and BrMC4 Tat displayed a diminished capacity to reactivate HIV transcription (**Fig. 4C**). Importantly, these differences were not attributable to cytotoxicity, as cell viability remained comparable across all transfection conditions.

Collectively, these findings demonstrate that microglia-derived HIV-1 strains exhibited lower activity in Tat-mediated transactivation of HIV, which further inhibited HIV transcription to enforce HIV latency in either T cells or microglia.

### Attenuated activities in HIV-1 transactivation by BrMC2 and BrMC4 Tat Variants are not due to impaired nuclear translocation

Previous studies have shown that the nuclear import of Tat is dependent on its basic domain, and mutations within this region can disrupt nuclear localization and function^44^. To determine whether impaired Tat nuclear translocation contributes to the reduced HIV-1 transcription observed in BrMC2 and BrMC4 Tat variants, we analyzed the subcellular distribution of each Tat variant. We extracted proteins from nuclear, cytoplasmic, and whole-cell fractions after transfection with Tat plasmids from different viral isolates and performed Western blot analysis. As expected, Tat proteins derived from pNL4.3 and HXB2 were slightly smaller than brain/MG-derived HIV Tat variants due to the absence of residues 87-101. We found that in 2D10 cells, cytoplasmic Tat expression was higher in BrMC2- and BrMC4-derived Tat proteins, while nuclear Tat remained constant except for YU-2 and JRCSF, compared to pNL4.3 or HXB2 (**Fig. 5A**). This was similarly observed in the HIV latently infected microglia model (**Fig. 5B**). Tat nuclear translocation remained intact in cells transfected with BrMC2/4 Tat plasmids. Thus, impaired transactivation by BrMC2 and BrMC4 Tat variants appears to result primarily from intrinsic defects in Tat function rather than from impaired nuclear translocation. Therefore, different from previous reports^44^, R52 may not be strictly required for Tat nuclear translocation.

**Figure 5.**
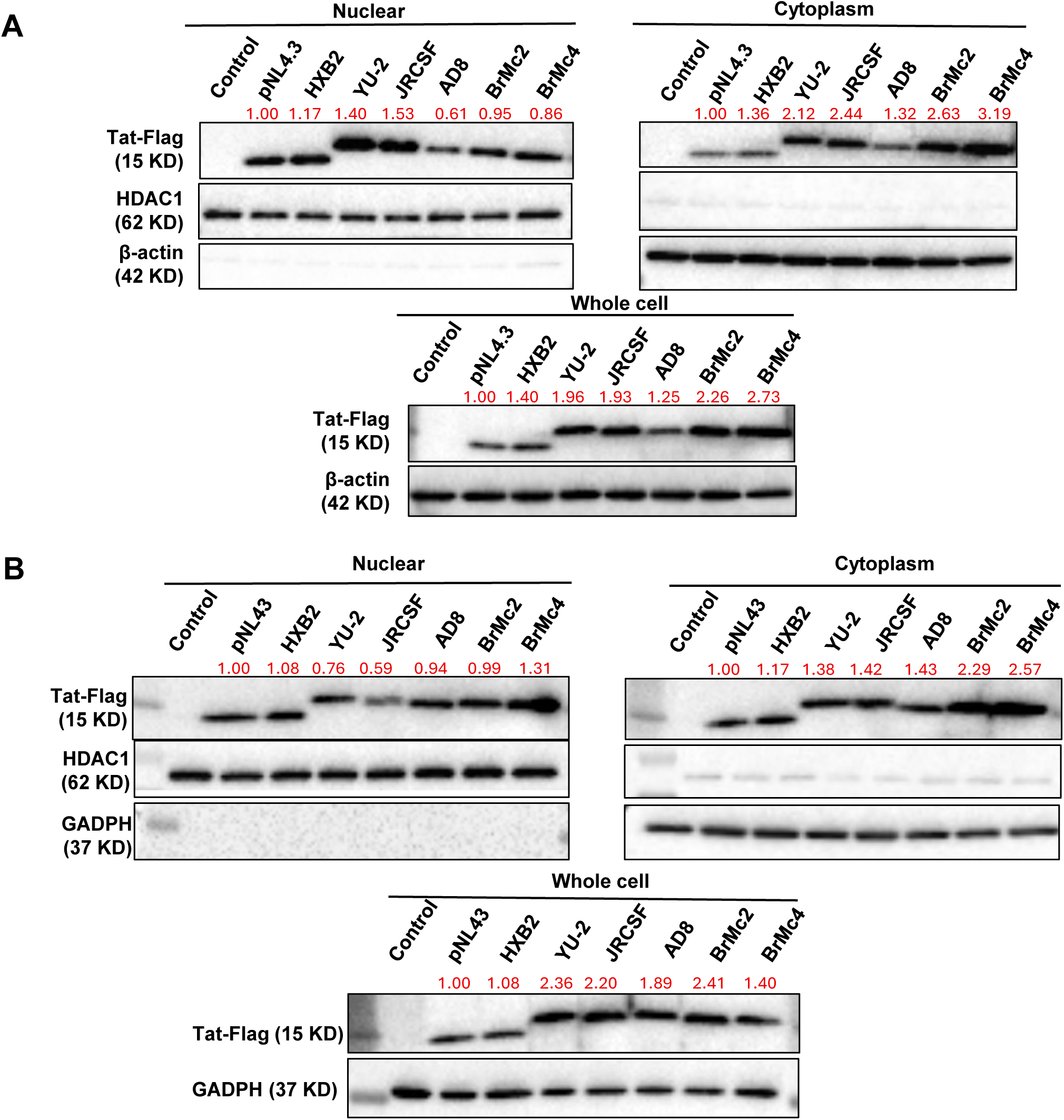
Subcellular localization of HIV-1 Tat protein variants. **(A-B)** Representative Western blot images of Tat protein in nuclear, cytoplasmic, and whole-cell fractions from 2D10 cells (A) and microglia (B) transfected with the indicated Tat-Flag plasmids. Densitometric analysis of the blot showing the nuclear and cytoplasmic expression of Tat protein (Normalized to pNL4.3-Tat). Due to a short HIV-1 Tat in pNL4.3, its molecular weight was slightly smaller than that of the other HIV-1 strains.

### Reverting the R52W Mutation Restores HIV-1 Transcription in BrMC2 and BrMC4 Tat Variants

Given the lower Tat transactivation activity of microglia-derived HIV-1 Tat independent of R52 on Tat nuclear translocation, we sought to understand whether the R52W mutation accounted for the attenuated Tat transactivation. The R52W mutation was identified as the sole amino acid substitution within the TAR-binding domain of both BrMC2 and BrMC4 Tat variants (**Fig. 2A**). As arginine at position 52 plays a critical role in mediating high-affinity with TAR RNA, we sought to determine whether this mutation impairs Tat-transactivation of HIV. To test this idea, we engineered revertant plasmids (BrMC2 W52R and BrMC4 W52R) that reverted tryptophan to arginine to restore the wild-type residue at position 52 (**Fig. 6A**). Upon transfection of the revertant into the 2D10 reporter cell line, we observed a marked increase in the percentage of GFP-positive cells relative to cells expressing the R52W mutants, reaching levels comparable to cells transfected with HXB2-derived Tat. These results indicate a significant enhancement in Tat-driven transcriptional activation (**Fig. 6B**). In TZM-bl cells, both revertant constructs (BrMC2 W52R and BrMC4 W52R) yielded significantly higher HIV transcription activity than their R52W mutant counterparts, reaching levels similar to the HXB2-Tat plasmids (**Fig. 6C**). Lastly, revertant constructs also restored Tat-driven HIV transcription in primary microglial cells (**Fig. 6D**). In all these models, transfections did not impact cell viability **(Supplementary Fig. 5)**. These results further underscore the critical role of arginine at position 52 in effective Tat-mediated transactivation.

**Figure 6.**
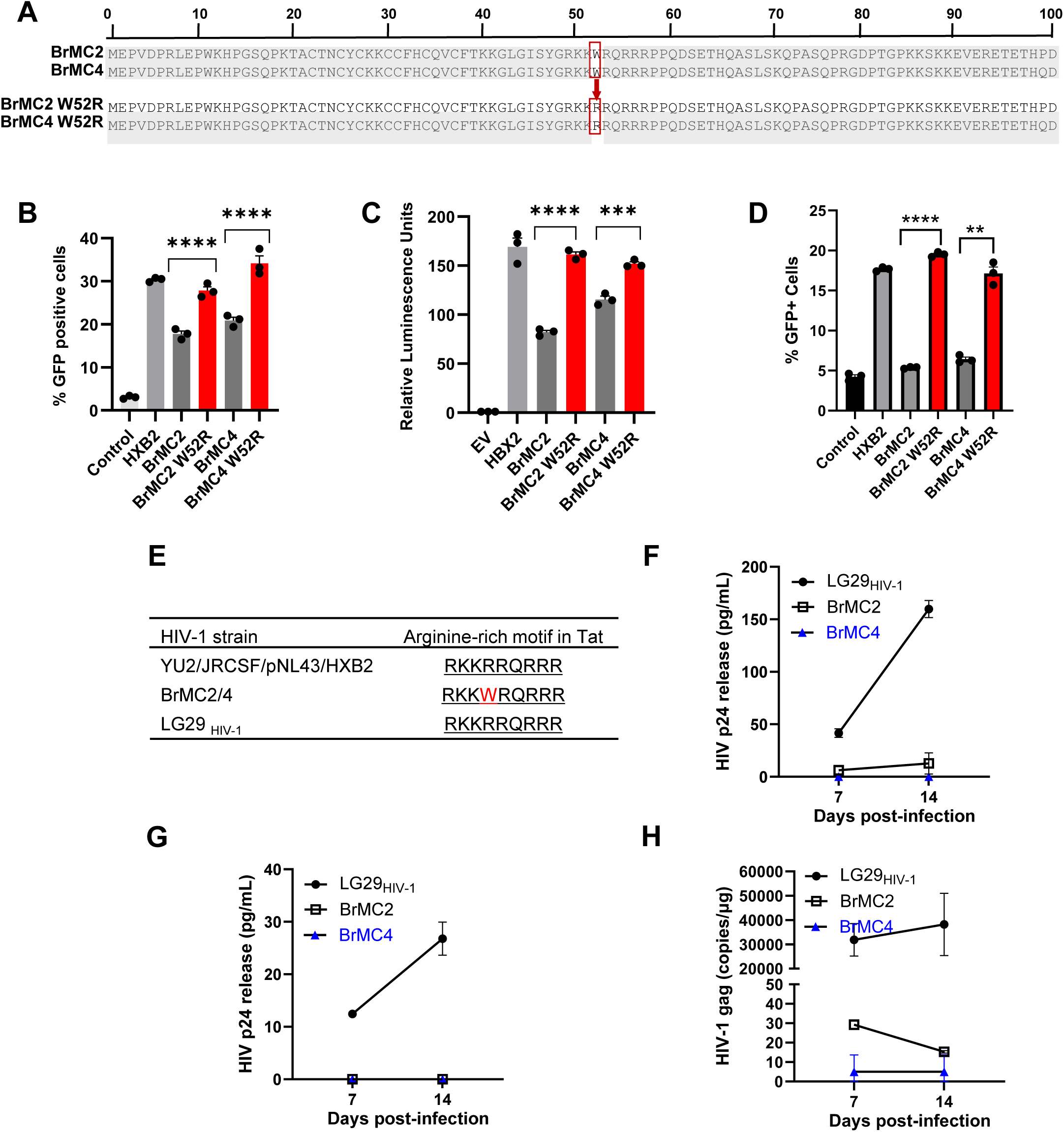
BrMC2 and 4 HIV-1 variants containing the R52W mutation attenuate HIV-1 infection in T cells and brain MG isolated from HIVneg donors. **(A)** Schematic diagram of the site-directed mutagenesis strategy used to introduce the W52R point mutation into the Tat gene of the BrMC2 and BrMC4 isolates. **(B)** and **(D)** Flow cytometry analysis of GFP expression in 2D10 (F) and microglial cells (G) 7 days post-transfection with W52 or W52R mutant Tat plasmids (2 µg). The geometric mean fluorescence intensity was used to assess Tat-dependent transactivation at the single-cell level. **(C)** W52R mutation restored Tat-driven HIV transcription. TZM-bl cells, which contain an integrated HIV-1 LTR-driven luciferase reporter, were transfected with plasmids expressing W52 (BrMC2/4) or W52R mutant Tat. Luciferase activity was measured to quantify LTR transactivation. **(E)** ARM sequences among different HIV-1 strains. **(F-H)** HIV-1 infection in the brain microglia and peripheral CD4^+^ T cells isolated from HIV negative donors. The same amount of HIV-1 was used during infection. Seven to fourteen days post-infection, the culture supernatants were collected, and HIV-1 replication in microglia was measured by p24 ELISA and/or ddPCR.

Lastly, we compared the ARM sequence of BrMC2/4 with that in LG29_HIV-1_, the replication-competent HIV-1 we recently recovered, sequenced, and characterized in the brain microglia ^1^, and noticed that the ARM sequence in LG29_HIV-1_ was intact, as in other well-known HIV-1 strains (**Fig. 6E**). Thus, we hypothesized that these sequence differences might contribute to distinct infection efficiencies between LG29_HIV-1_ and microglia-derived BrMC2/4. To this end, we isolated microglia from HIV-negative donors as in **Figure 1A**. We found that infection of HIV-negative microglia by R52W-mutant BrMC2 and BrMC4 HIV-1 was markedly less efficient than by ARM-intact LG29_HIV-1_ (**Fig. 6F**), where p24 in microglia infected with BrMC4 was undetectable. Similar results were observed in primary CD4^+^ T cells, where supernatant p24 in BrMC2 or BrMC4-infected cells was almost undetectable by ELISA (**Fig. 6G**), unless measured by a highly sensitive RT-qPCR assay (**Fig. 6H**). Together, these observations support a crucial role of a naturally occurring R52W Tat mutation in attenuation of HIV transcription and replication in T cells and non-T cells.

### HIV-1 recovered from the microglia is found in the peripheral compartment long before ART interruption

Despite HIV outgrowth in several wells, only two replication-competent microglia-derived HIV-1 clones could be recovered and sequenced. Notably, both isolates contained an identical Tat R52W substitution. To investigate the origin and tissue distribution of this variant, we analyzed HIV-1 DNA and RNA sequences obtained from multiple anatomical compartments of the same donor (**Fig. 7A**). The R52W substitution was absent from HIV-1 DNA sequences recovered from the basal ganglia but was detected in several peripheral compartments, including the spleen and duodenum. Longitudinal analyses further revealed that R52W was already present in peripheral viral populations prior to death, including plasma HIV RNA collected during viral rebound (14 February 2022, ∼300 days prior to death) and PBMC-associated HIV sequences obtained approximately one year earlier (13 December 2021, 1 day prior to death). At autopsy (14 December 2022), the same substitution was identified in viral sequences recovered from the spleen and in the microglia-derived outgrowth viruses BrMC2 and BrMC4 (**Fig. 7B**).

**Figure 7.**
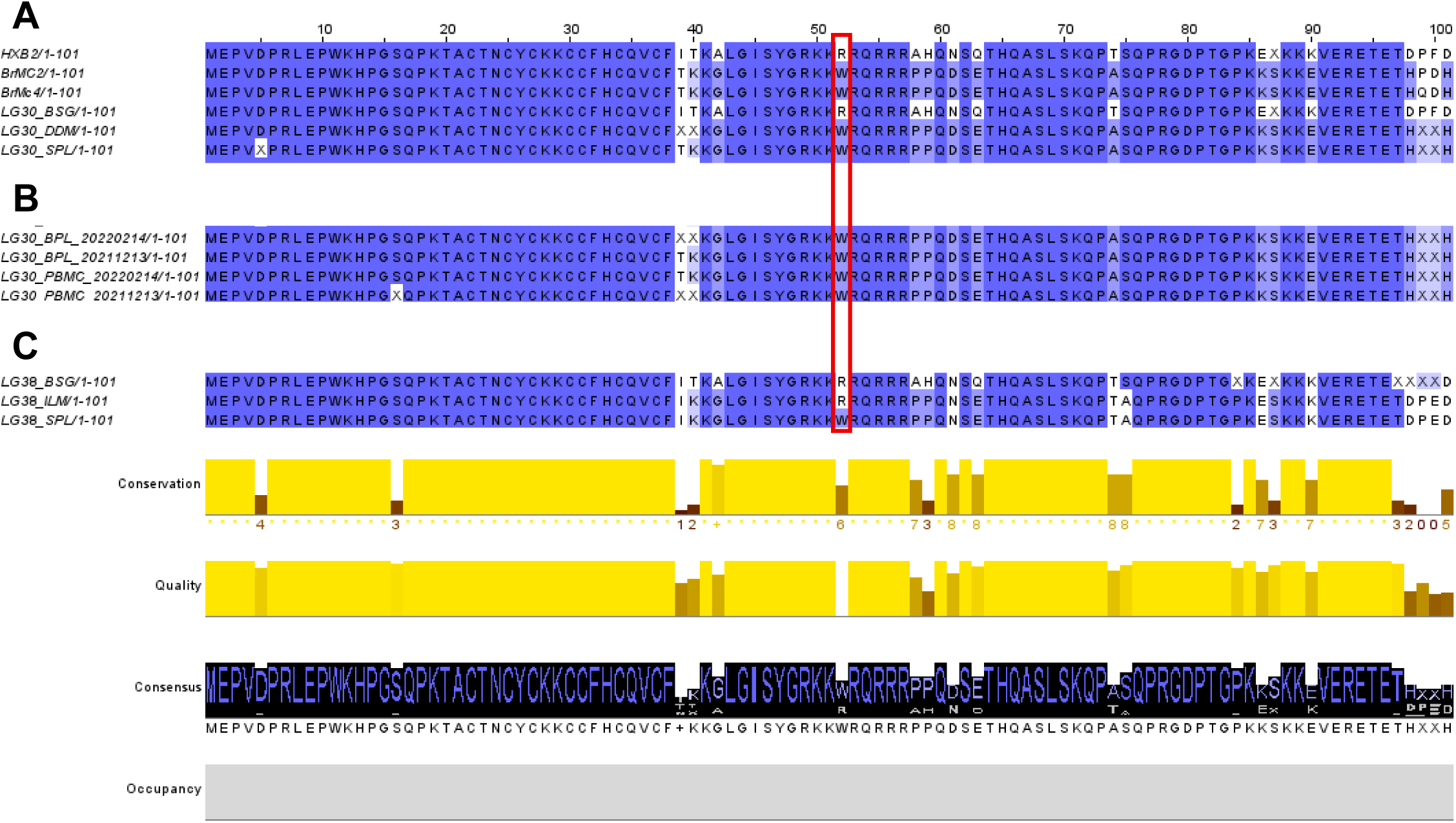
Sequence alignment of Tat derived from outgrown HIV-1 in microglia and HIV-1 from the basal ganglia, as well as peripheral compartments. **(A)** Amino acid sequence alignment of the HIV-1 Tat proteins derived from outgrowth HIV-1 RNA sequences from microglia and blood plasma (BPL), with DNA sequences from basal ganglia (BSG), dedendum (DDW), PBMCs, or spleen tissues of the same PWH, where HXB2 sequence serves as an HIV-1 reference genome. **(B)** The Tat gene was longitudinally sequenced during infection, ART treatment, and until one day before death. **(C)** The same R52W Tat mutation was found in the spleen of another PWH, LG38, on ART.

To determine whether the Tat R52W variant is present in other PWH, we analyzed genomic DNA from multiple tissues, including the basal ganglia, ileum, and spleen, from an independent donor (LG38) on suppressive ART in the same cohort. Notably, the R52W mutation was detected in HIV-1 Tat sequences from the spleen but not from the brain or gut (**Fig. 7C**), suggesting that this variant may emerge repeatedly in distinct anatomical compartments among PWH on suppressive ART.

### Tat residue 52 is highly conserved across global HIV-1 isolates but exhibits a naturally occurring W52 polymorphism

To further determine whether the R52W substitution represents a naturally occurring HIV-1 polymorphism, we analyzed 3,440 unique Tat protein sequences recovered from plasma and/or PBMCs of PWH obtained from the Los Alamos National Laboratory HIV Sequence Database. Sequence alignments were generated relative to the HXB2 reference sequence, and amino acid frequencies were quantified at position 52, which lies within the Tat arginine-rich motif (ARM) responsible for TAR RNA recognition.

Residue conservation analysis revealed that arginine at position 52 is highly conserved across global HIV-1 isolates, occurring in 3,353 of 3,440 sequences (97.47%) (**Table 2**). In contrast, tryptophan at this position was identified in only 73 sequences (2.12%), establishing the R52W substitution as an uncommon but naturally occurring HIV-1 variant. Other amino acids were observed only rarely at this position, including glutamine (0.17%), lysine (0.03%), glycine (0.03%), ambiguous residues (0.09%), and alignment gaps (0.09%).

**Table 2.** Global conservation analysis of HIV-1 Tat residue 52 across 3,440 unique HIV-1 isolates. Residue frequencies at position 52 of HIV-1 Tat were determined from a multiple sequence alignment of 3,440 unique Tat protein sequences obtained from the Los Alamos National Laboratory HIV Sequence Database. Residue numbering is based on the HXB2 reference sequence (accession K03455). Position 52 lies within the Tat arginine-rich motif (ARM), which mediates TAR RNA recognition and transcriptional activation.

| Residue at Position 52 | Number of Sequences | Frequency (%) |
| --- | --- | --- |
| R | 3,353 | 97.47 |
| W | 73 | 2.12 |
| Q | 6 | 0.17 |
| X (Ambiguous residue) | 3 | 0.09 |
| Gap (-) | 3 | 0.09 |
| K | 1 | 0.03 |
| G | 1 | 0.03 |
| Total | 3,440 | 100.00 |

Residue numbering corresponds to the HIV-1 HXB2 Tat reference sequence (B.FR.1983.HXB2LAIIBBRU.K03455). Ambiguous residues (X) represent positions for which the deposited sequence did not allow unambiguous amino acid assignment.

Examination of available clinical metadata revealed that 69 of 73 (94.5%) R52W-containing sequences were derived from blood or PBMC samples obtained from ART-naïve individuals. Notably, the remaining four sequences originated from individuals receiving suppressive ART (**Table 1**)^47, 48, 49^, and all four carried the R52W substitution. Among these cases, one individual maintained virologic control following analytical treatment interruption (ATI)^47^.

To determine whether the W52 variant was restricted to a particular HIV-1 lineage, we next examined its subtype distribution (**Table 3**). Subtype B constituted the largest dataset (n = 2,260), followed by subtype C (n = 994), subtype D (n = 174), and subtype A (n = 12). R52 remained highly conserved across all major subtypes, occurring in 97.61% of subtype B sequences, 96.88% of subtype C sequences, 98.85% of subtype D sequences, and 100% of subtype A sequences. Importantly, the W52 substitution was detected in multiple independent viral lineages, including subtype B (45 sequences; 1.99%), subtype C (27 sequences; 2.72%), and subtype D (1 sequence; 0.57%). No W52 variants were observed among the limited number of subtype A sequences analyzed. These findings indicate that the R52W substitution is not restricted to a single HIV-1 subtype but instead represents a low-frequency naturally occurring polymorphism distributed across genetically distinct viral lineages.

**Table 3.**
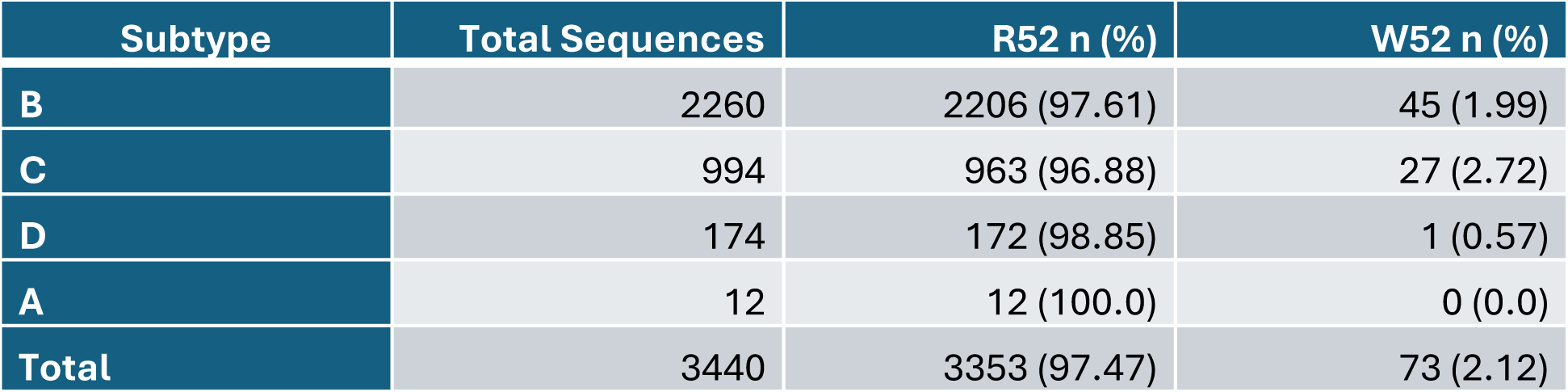
Subtype-specific analysis of HIV-1 Tat residue 52.

Sequence logo analysis of the Tat ARM region further revealed strong evolutionary conservation of the basic residues required for TAR recognition, consistent with the central role of this domain in HIV transcriptional activation (**Figure 8**). Among these residues, R52 exhibited exceptionally high conservation across global HIV-1 isolates, suggesting strong evolutionary constraint at this position. Substitution of R52 with tryptophan therefore represents a non-conservative change that simultaneously removes a positively charged side chain and introduces a bulky hydrophobic aromatic residue within the RNA-binding interface.

**Figure 8.**
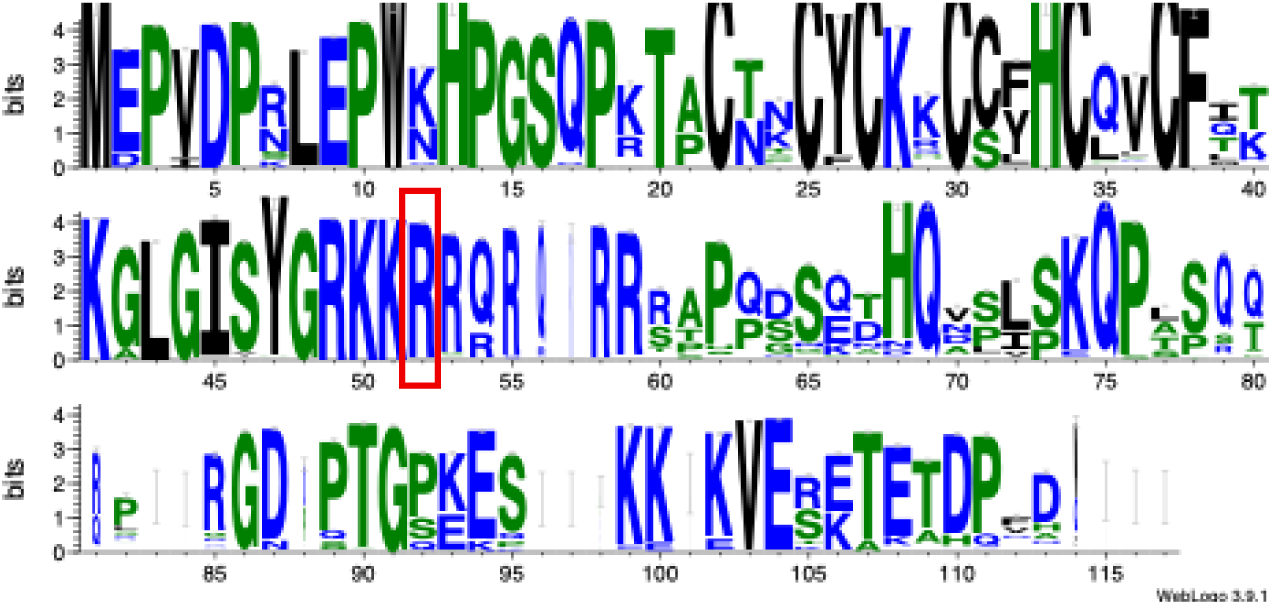
Sequence conservation analysis of HIV-1 Tat reveals strong evolutionary constraint at residue 52 within the ARM. Sequence logo representation generated from a multiple sequence alignment of 3,440 unique HIV-1 Tat protein sequences obtained from the Los Alamos National Laboratory HIV Sequence Database using a one-sequence-per-patient filtering strategy. Residue numbering is based on the HIV-1 HXB2 reference sequence. The height of each amino acid letter is proportional to its frequency at a given position, and the total stack height reflects sequence conservation. Red rectangle: R52 residue in HIV Tat.

Collectively, these analyses establish that R52 is a highly conserved residue within the Tat ARM, whereas the W52 variant represents a rare but naturally occurring HIV-1 polymorphism detected across multiple viral subtypes and anatomical compartments, including in individuals receiving suppressive ART. When considered together with its association with impaired Tat–TAR interactions and reduced transactivation activity, these findings suggest that R52W may define a distinct functional Tat variant with potential relevance to viral persistence and reservoir biology.

## Discussion

Previously, we reported two near full-length replication-competent HIV-1 isolates from brain microglia of a single PWH^1^. In the present study, we extend these findings by characterizing two full-length HIV-1 genomes from brain microglia of an additional donor in the Last Gift cohort, with a focus on the Tat–TAR regulatory interface.

We identify a naturally occurring R52W substitution within the brain microglia-derived Tat arginine-rich motif that is associated with attenuated viral transcriptional activity. Of note, this R52W variant, detected in brain microglia and peripheral tissues on ART, was also observed in the spleen in an independent individual receiving suppressive ART. Expanding these findings, across a dataset of 3,440 clinical HIV-1 isolates, the R52W substitution was identified in 2.12% of sequences, including multiple ART-suppressed individuals and a therapy controller. The recurrence of this variant across genetically diverse viral lineages supports its presence in ART-suppressed populations rather than a compartment-specific or isolated event.

The absence of R52W in basal ganglia–associated HIV DNA, together with its longitudinal detection in peripheral compartments, is consistent with a peripheral origin of this variant followed by dissemination into the CNS. Although the directionality of viral trafficking cannot be established from these data alone, the temporal and anatomical distribution of R52W supports the existence of a persistent peripheral viral lineage capable of contributing to the CNS reservoir. Future studies combining longitudinal phylogenetic reconstruction, single-genome sequencing across anatomical compartments, and integration site analyses will be required to determine the evolutionary origin of the R52W lineage, define the directionality of viral migration between peripheral tissues and the CNS, and establish whether this variant confers a selective advantage within specific tissue reservoirs.

Despite conservation of the TAR RNA structure, the R52W substitution selectively disrupts Tat function, revealing a mechanism by which variation in a highly conserved regulatory protein can modulate viral transcription without altering the RNA target element. The Tat ARM (residues 49–57; 49-RKKRRQRRR-57) is essential for TAR binding, nuclear localization, and Tat-mediated transactivation, and is among the most functionally constrained regions of the viral genome^17, 50^. Consistent with prior mutagenesis studies showing that substitutions at R52 substantially impair Tat activity, our structural modeling indicates that R52W decreases Tat–TAR stability and reduces hydrogen bonding interactions

Functional assays confirmed that R52W is sufficient to markedly reduce Tat-mediated transcriptional activity. A genetic reversion experiment (W52R) restored transcriptional function, establishing a direct causal role for residue 52 in maintaining Tat activity. These findings demonstrate that single naturally occurring substitutions within the ARM can significantly alter Tat function through disruption of RNA–protein interaction dynamics.

Functionally, the R52W substitution attenuates Tat-mediated transcription, resulting in reduced viral gene expression. This transcriptionally attenuated phenotype is consistent with a state potentially compatible with persistence in long-lived cellular reservoirs. Importantly, recurrence of this variant in additional PWH on ART, including a therapy controller, supports its presence under conditions of viral suppression. While these observations do not establish a causal role for R52W in promoting persistence, they suggest that transcriptionally attenuated viral variants may contribute to functional heterogeneity within the reservoir.

In addition to the ARM mutation, additional substitutions observed in the Proline-rich, Cysteine-rich, and Core domains of Tat in these isolates may further contribute to impaired transactivation by affecting Cyclin T1/CDK9 binding and overall protein stability^16^, although their individual contributions require further investigation.

Several limitations should be noted. The number of microglia-derived full-length viral genomes remains limited, and thus, the full extent of HIV genetic diversity within CNS myeloid reservoirs is not fully captured. In addition, functional analyses were performed in vitro and do not directly measure the impact of these variants on reservoir dynamics in vivo. Future studies using more physiologically relevant models, including ex vivo tissue systems or humanized animal models, will be required to define their role in viral persistence. Also, the initial Tat–TAR complexes were generated using AlphaFold3 rather than experimental structures. While numerous structures of HIV-1 Tat–TAR complexes have been reported, these correspond primarily to laboratory strains and do not capture the sequence diversity of the patient-derived variants investigated here. AlphaFold3 enabled construction of variant-specific complexes using a consistent modeling strategy across all systems. The predicted models reproduced the conserved Tat–TAR interaction topology and served as starting conformations that were subsequently refined during 1-μs molecular dynamics simulations. Moreover, an independent simulation performed using an AMBER ff19SB/OL3 force-field combination yielded similar interaction patterns and conformational trends (**Supplementary Fig. 4**), supporting the robustness of our conclusions.

In summary, this study links naturally occurring variation in HIV-1 Tat to a defined molecular mechanism regulating transcriptional activity and supports the concept that functional diversity in viral regulatory proteins contributes to persistent HIV infection under suppressive ART.

## Data Availability Statement

All relevant data are within the manuscript and its Supporting Information files.

## Contributions

GJ conceived the study. SG provides brain tissues after rapid research autopsies. YT performed the approaches to recover HIV-1 from the brain microglia. AC and HC analyzed HIV-1 genome sequences. HC validated their functions in latency models in vitro. DMM provides necessary resource support at the UNC HIV Center. VRC modeled HIV Tat–TAR complexes and performed trajectory and structural analyses. Working with VRC, NJ modeled Tat/TAR-1 interaction with thousands of HIV genomes from PWH. HC, GJ, and VRC revised and edited multiple versions. GJ finalized the manuscript for submission. All authors approved the final manuscript.

## Acknowledgements and Financial Disclosure Statement

We thank Dr. Katharine Barr for sequencing the HIV-1 genomes and Dr. Jonathan Karn for providing us with the 2D10 model of HIV latency. This work was supported by R21MH128034, R01MH136852, R01MH1394460-01A1, R21AI167709, R01AI186609, UM1AI164567, and the Collaborative Development Program at B-HIVE (U54AI170855) to GJ, as well as by UNC Cancer Center Core Support Grant P30CA016086 to VRC.

---

**Supplementary Figure 1.**
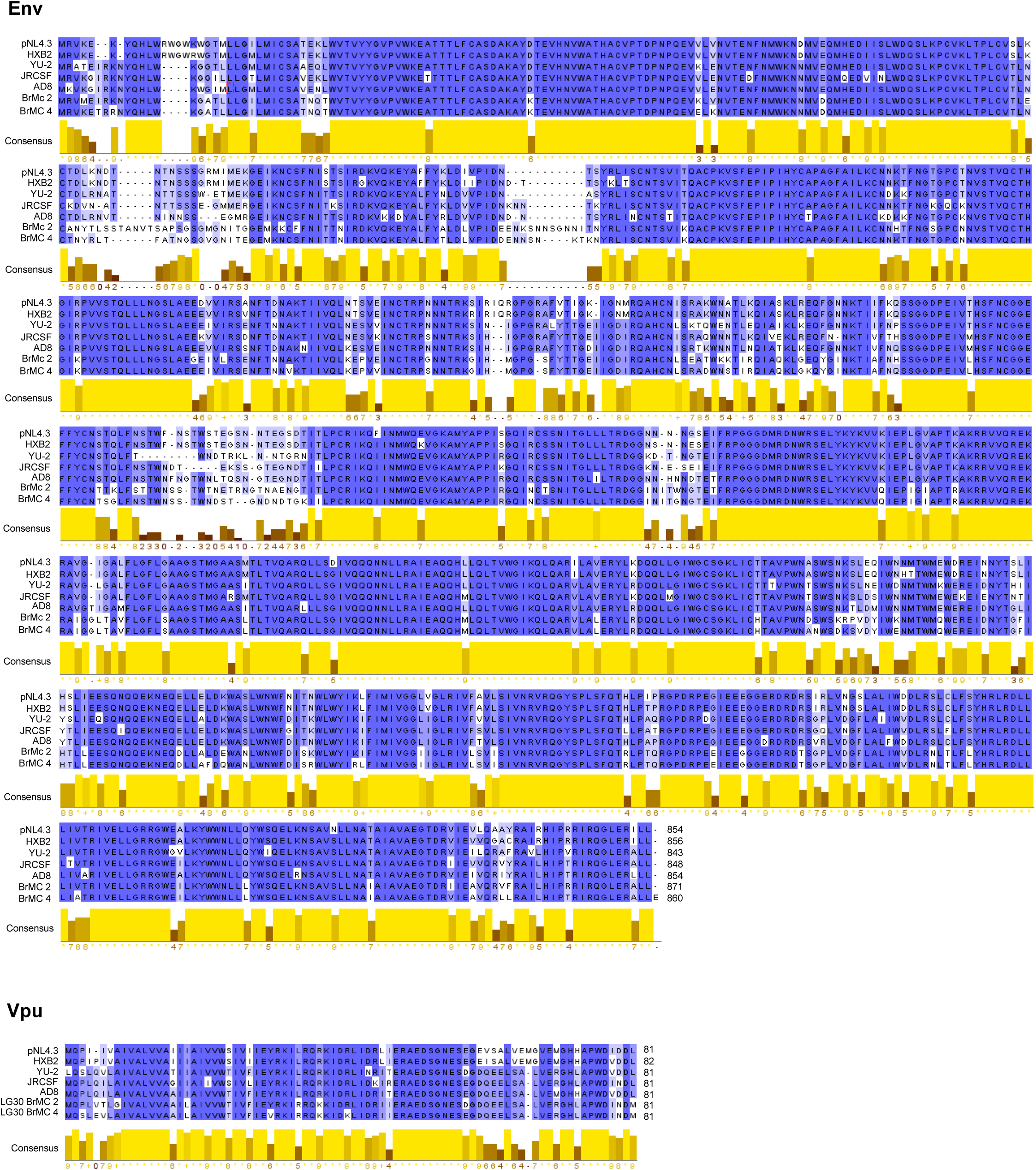
Protein alignment of HIV-1 Env and Vpu. Amino acid sequence alignments of HIV-1 Env and Vpu, comparing viral sequences isolated from brain microglia (BrMC2 and BrMC4) with commonly referenced HIV-1 strains (pNL4.3, HXB2, YU-2, JRCSF, AD8).

**Supplementary Figure 2.**
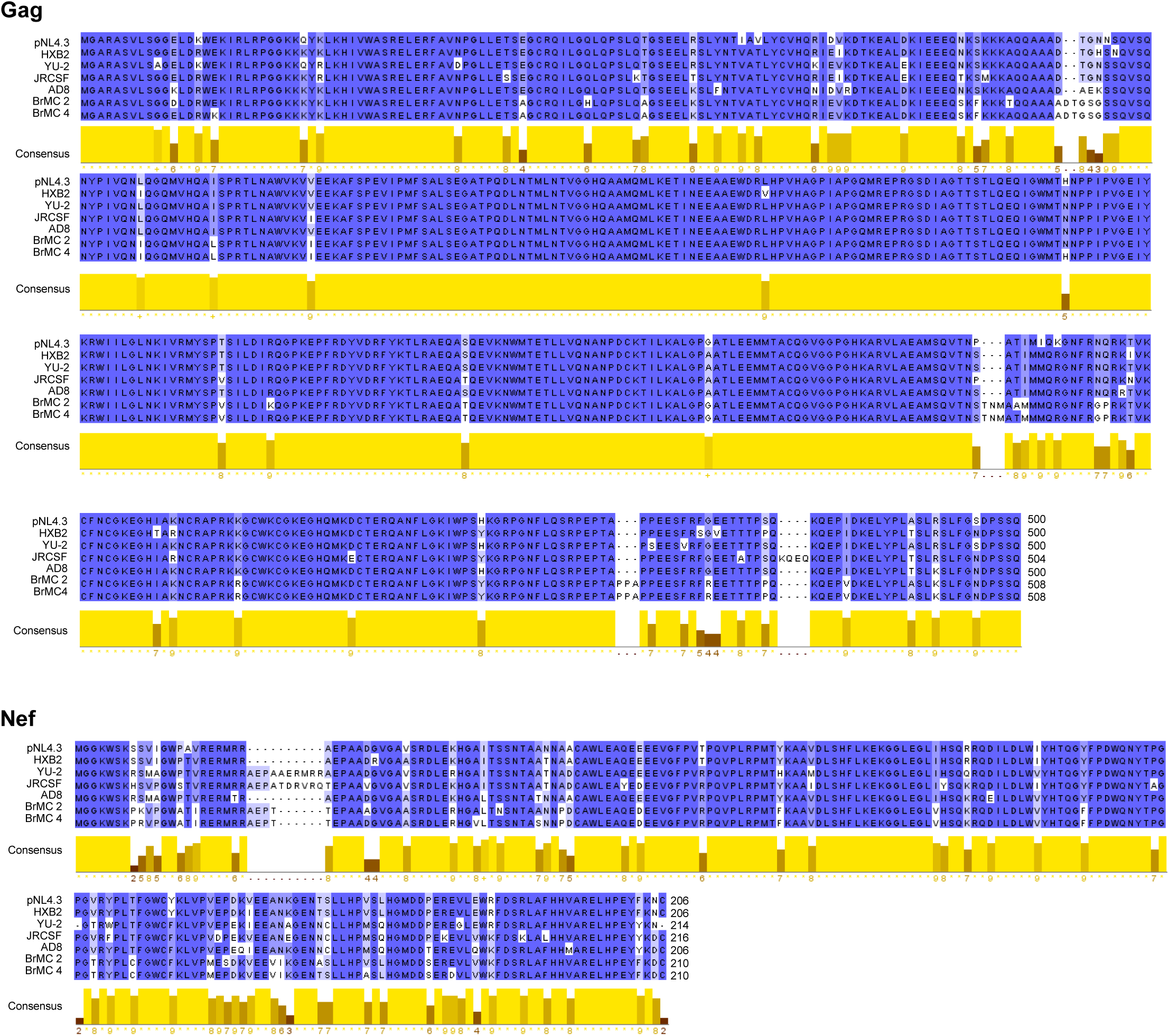
Protein alignment of HIV-1 Gag and Nef. Amino acid sequence alignments of HIV-1 Gag and Nef comparing viral sequences isolated from brain microglia (BrMC2 and BrMC4) with commonly referenced HIV-1 strains (pNL4.3, HXB2, YU-2, JRCSF, AD8).

**Supplementary Figure 3.**
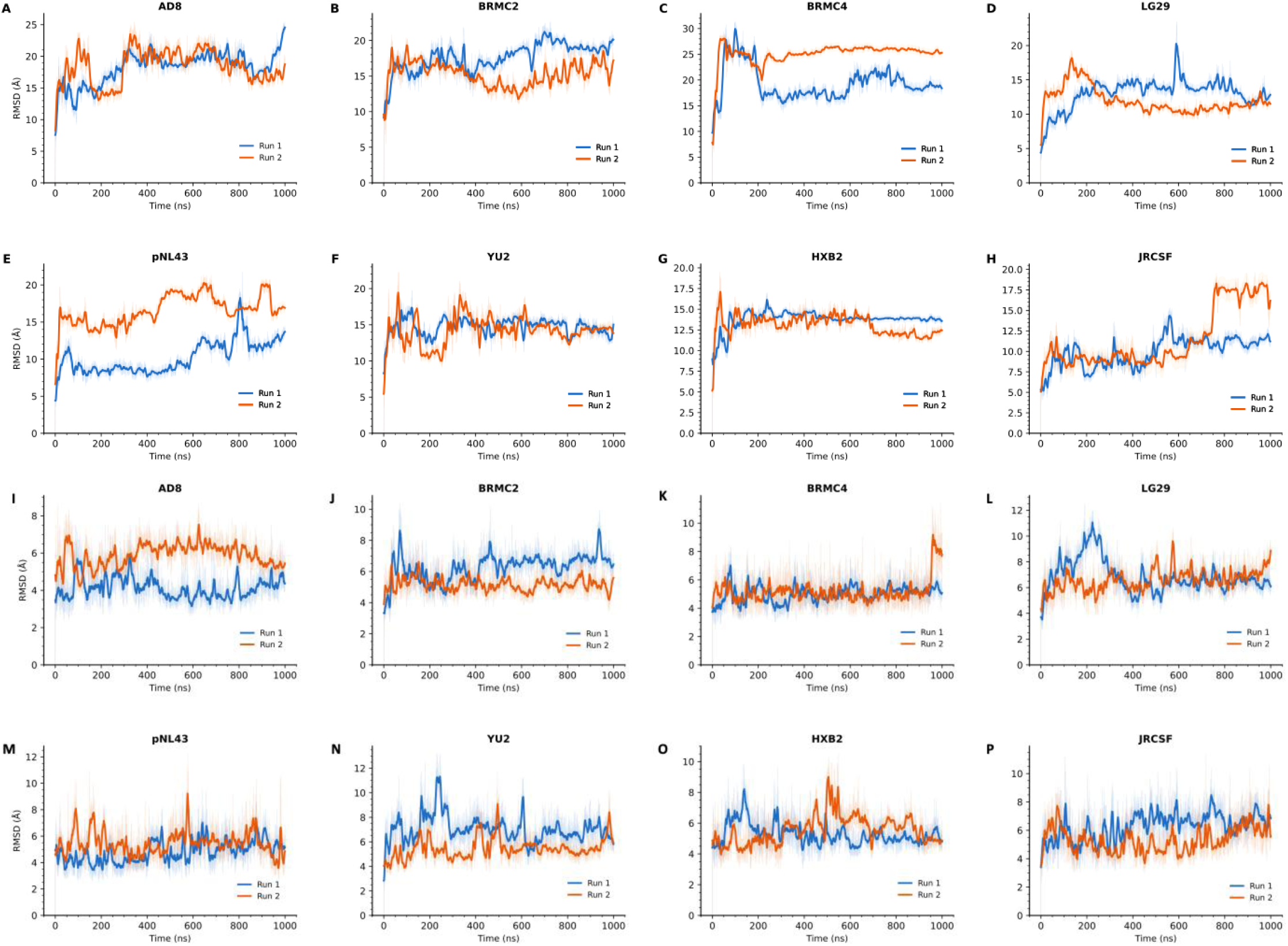
Replica consistency of Tat-TAR structural dynamics across all HIV isolates. Root mean square deviation (RMSD) was calculated from two independent MD simulations (Run 1, blue; Run 2, red) for all eight HIV Tat–TAR complexes. (A–H) Tat backbone Cα RMSD and (I–P) TAR RNA RMSD for AD8, BRMC2, BRMC4, LG29, pNL43, YU2, HXB2, and JRCSF, respectively.

**Supplementary Figure 4.**
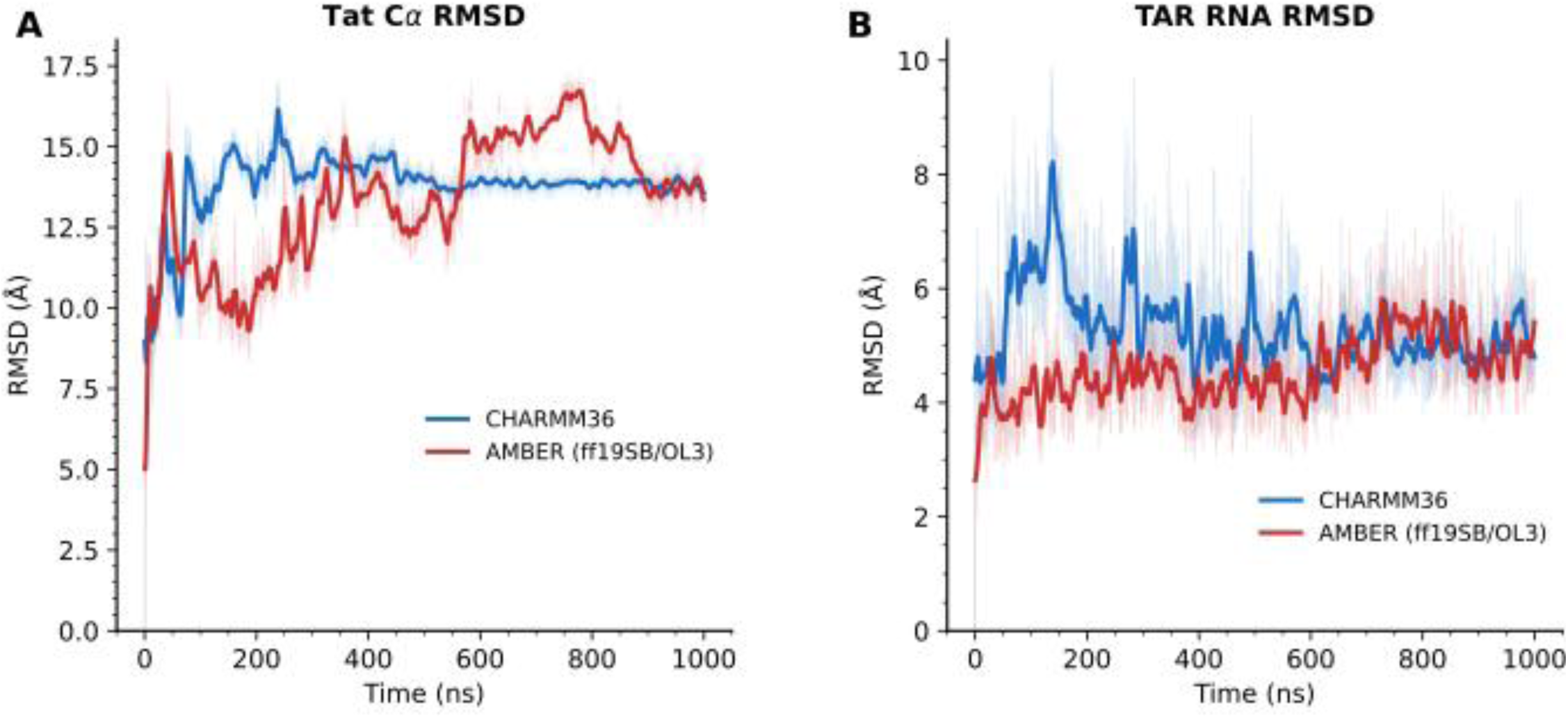
Force field comparison of Tat–TAR complex structural stability assessed by RMSD analysis. RMSD of the HIV HXB2 Tat–TAR complex was calculated from MD simulations performed with two independent force field combinations: CHARMM36 (blue) and AMBER (ff19SB for protein/OL3 for RNA; red). (A) Backbone RMSD of Tat protein Cα atoms, calculated after least-squares fitting to the initial Cα coordinates. (B) RMSD of TAR RNA heavy atoms, calculated after least-squares fitting to the initial TAR coordinates.

**Supplementary Figure 5.**
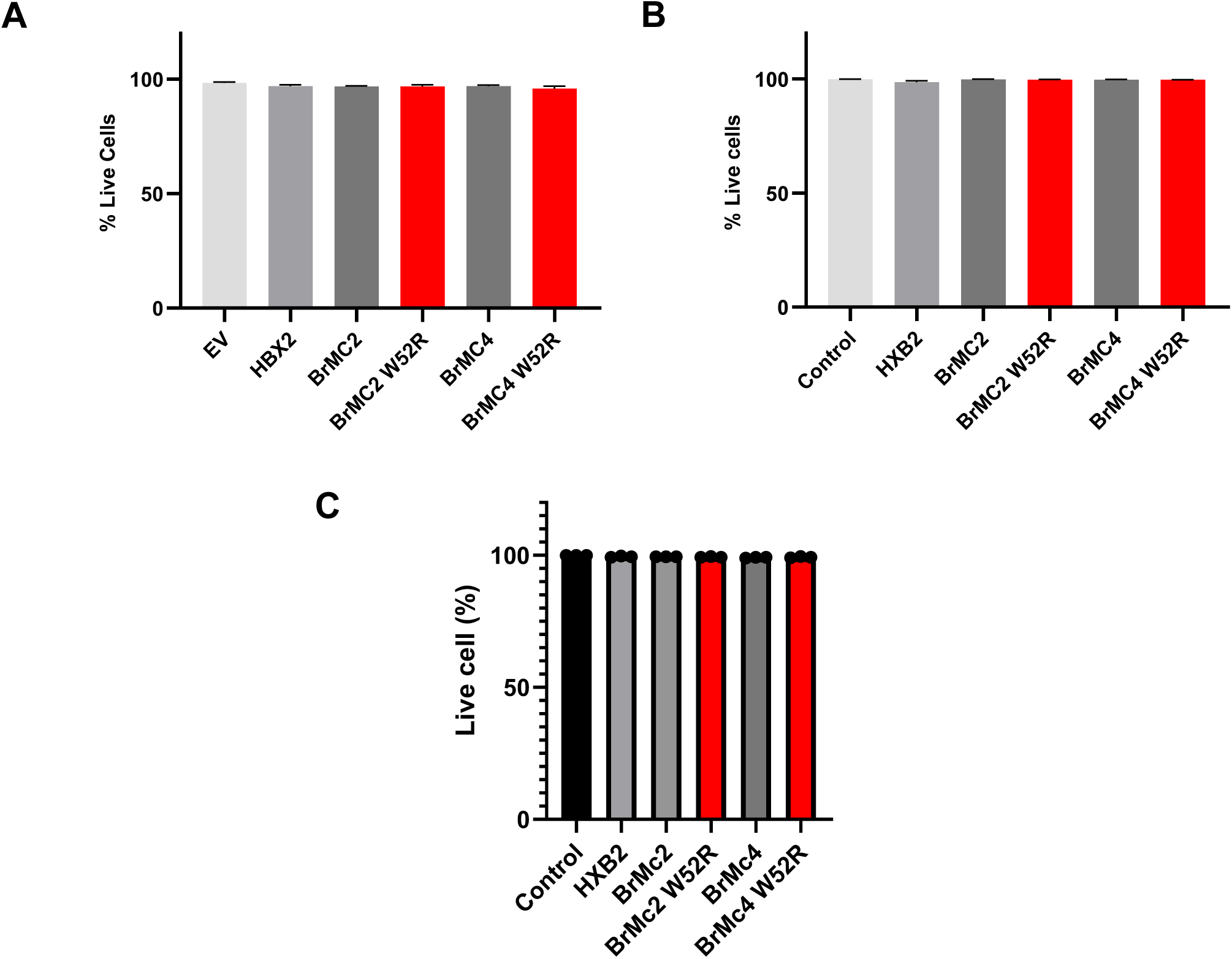
The R52W mutation impairs Tat-mediated transactivation in brain-derived HIV-1 isolates without impact on cellular viability. Flow cytometry analysis of cellular viability of 2D10 cells **(A)**, TZM-bl cells **(B)**, and microglial cells **(C)** 7 days post-transfection with W52 or W52R mutant Tat plasmids.

**Supplementary Table 1.** AlphaFold3 confidence metrics for HIV-1 Tat–TAR complexes across viral variants.

| Variant | Ranking Score | pTM | iPTM | Minimum Inter-chain PAE (Å) | Steric Clash | Model Used |
| --- | --- | --- | --- | --- | --- | --- |
| pNL4-3 | 0.59 | 0.38 | 0.22 | 4.95 | None | Model 0 |
| HXB2 | 0.59 | 0.36 | 0.22 | 4.76 | None | Model 0 |
| JR-CSF | 0.58 | 0.38 | 0.23 | 5.14 | None | Model 0 |
| AD8 | 0.57 | 0.33 | 0.17 | 4.95 | None | Model 0 |
| LG29 | 0.57 | 0.33 | 0.18 | 4.95 | None | Model 0 |
| BRMC2 | 0.54 | 0.32 | 0.15 | 6.54 | None | Model 0 |
| BRMC4 | 0.55 | 0.33 | 0.17 | 5.29 | None | Model 0 |
| YU-2 | 0.68 | 0.41 | 0.25 | 4.74 | None | Model 0 |

